# The microbiome as a biosensor: functional profiles elucidate hidden stress in hosts

**DOI:** 10.1101/752261

**Authors:** Avihai Zolti, Stefan J. Green, Noa Sela, Yitzhak Hadar, Dror Minz

## Abstract

Microbial communities are highly responsive to environmental cues, and both their structure and activity can be altered in response to changing conditions. We hypothesized that host-associated microbial communities, particularly those colonizing host surfaces, can serve as *in situ* sensors to reveal environmental conditions experienced by both microorganisms and the host. For a proof-of-concept, we studied a model plant-soil system and employed a non-deterministic gene-centric approach. A holistic analysis was performed using plants of two species and irrigation with water of low quality to induce host stress. Our analyses examined the genetic potential (DNA) and gene expression patterns (RNA) of plant-associated microbial communities, as well as transcriptional profiling of host plants. Transcriptional analysis of plants irrigated with treated wastewater revealed significant enrichment of general stress-associated root transcripts relative to plants irrigated with fresh water. Metagenomic analysis of root-associated microbial communities in treated wastewater-irrigated plants, however, revealed enrichment of more specific stress-associated genes relating to high levels of salt, high pH and lower levels of oxygen. Meta-analysis of these differentially abundant genes obtained from other metagenome studies provided evidence of the link between environmental factors such as pH and oxygen and these genes. Analysis of microbial transcriptional response demonstrated that enriched gene content was actively expressed, which implies contemporary response to elevated levels of pH and salt. We demonstrate here that microbial profiling can elucidate stress signals that cannot be observed even through interrogation of host transcriptome, leading to an alternate mechanism for evaluating *in situ* conditions experienced by host organisms.

**Significance Statement:** This study examines the potential for microbial communities to provide insight into stresses experienced by their eukaryotic host organisms, through profiling of metagenomes and metatranscriptomes. Our study uses plant host-associated microorganisms as an *in vivo* and *in situ* microsensor to identify environmental stresses experienced by the microbial community and by the plant. Transcriptionally active host-associated microbial communities are responsive in a highly specific manner to environmental conditions. Conversely, host transcriptome sequencing provides only a very general stress response. This study is a proof-of-concept for the use of microbial communities as microsensors, with a great potential for interrogation of a wide range of host systems.

## Introduction

Advances in sequencing have propelled the field of microbiology and shifted focus from analysis of microbial isolates or low diversity ecosystems to analysis of environments with highly diverse microbial communities. Global surveys of microbial community structure have been conducted in a wide range of natural environments (1, 2, 3), also reviewed in (4), and many of these studies have focused on host-associated microbiomes. Such host-associated microbial environments include plant-associated communities (5, 6, 7, 8, 9) though the greatest effort has been placed on the human-associated microbial communities (10, 11, 12, 13). Studies examining plant host-associated microbial communities have focused on soil microorganisms that are enriched in the rhizosphere-the soil surrounding and affected by the roots (8). In the rhizosphere, soil type (6, 8, 9) and plant host type (14, 15) have been identified as the main forces determining rhizosphere and root microbiomes. The selection of rhizosphere-competent organisms from soil has been well established, with specific plants and different growth stages of plants each selecting for different microbial communities from among the high diversity of microorganisms in soil (16, 17). Rhizosphere microorganisms are further enriched to form sub-populations colonizing root surface (9), as plants shape the soil-plant continuum in a gradient-depended manner (6, 8, 9) mainly through carbon flux to the root environment (18). Functional profiling of microbial communities associated with different plants has demonstrated that these microbiomes differ in their metabolic activities and has suggested the presence of niche conditions associated with a wide range of factors, of which oxygen concentration is one (19). More broadly, factors influencing plant root microbiome include geographic location (5, 9), plant developmental stage (15, 20, 21), nutrient (*e.g.,* N or P) availability (7, 22) and redox status (23). Numerous agricultural practices, which modify many of the above, have been shown to have an impact on root microbiome. These include fertilization (24), compost amendment (25) and irrigation with water of lower quality (26). Each of these practices alters a wide range of environmental variables, thus confounding the ability to identify the most consequential abiotic factor influencing the plant system and modifying the root microbiome.

Microorganisms sense minor changes in environmental conditions and respond rapidly through transcriptional changes, as well as through microbial amplification-the dynamic modification of the abundance of microbial taxa; these changes occur on a time-scale that is much shorter than for the host (27). Thus, interrogation of the microbial community may be used as a means to understand environmental conditions on short to long time scales, as well as small to large physical scale. In this study, the root surface is used as a model to test the hypothesis that microorganisms can be sensitive *in situ* detectors of environmental conditions. The root zone has a number of favorable features for such interrogation, including: (a) the presence of a high percentage of microorganisms that are transcriptionally active; (b) high microbial competition for access to root exudates – and therefore likely rapid turn-over if environmental conditions change; and (c) access to high microbial diversity in the soil. Thus, both the composition of the root-associated microbial community and the transcriptional activity of the microbial community can be informative regarding root environmental conditions.

In this study, we use the root surface microbiome functional response as a micro-sensor to identify stresses imposed by irrigation with water of lower quality, such as treated wastewater (TWW). Our study employs the basic assumptions that the microbiome inhabiting the root surface is exposed to the same environmental conditions as its host, and that the response of the microbiome (*i.e.*, alteration of community structure, associated gene abundance and transcriptional profiles) to stress can identify the specific stress or stresses in the root environment. To examine these assumptions and our general hypothesis, we performed deep DNA and RNA sequencing of plant roots grown in soil irrigated with fresh water (FW) or TWW. Plant transcriptional profiling was examined together with microbial community taxonomic and functional gene content characterization (shotgun metagenome sequencing) and microbial transcriptional profiling (shotgun metatranscriptome sequencing). We observed that the host responded to TWW irrigation in a highly general manner, whereas the microbial response was specific to stresses present in TWW, including elevated salinity and elevated pH.

## Results

Here, we assess the use of root-associated microorganisms as an indicator tool to reveal environmental conditions and stresses affecting plant hosts at a micro-scale. Our model system was a long-term anthropogenic disturbance caused by soil irrigation with water of lower quality (*i.e.,* treated wastewater, TWW) as compared to irrigation with fresh water (FW). We characterized plant-host response and root microbiome composition and response using deep sequencing of RNA and DNA extracted from roots. Shotgun metagenomic (DNA-based) and metatranscriptomic (RNA-based) analyses were performed on root systems from two host plant types (tomato and lettuce) across two consecutive years and with two different water treatments (FW or TWW), to describe taxonomic shifts and functional responses associated with long-term root irrigation with water of differing quality. For shotgun metagenome analyses, 25-49 million DNA sequences (paired-end) were generated per sample (**supplementary Table S1**). In tomato roots, 13 to 28% of the reads mapped to the host genome, with the remaining reads were attributed to the microbiome. In lettuce, 55-69% mapped to the host with the remaining reads attributed to the microbiome. A *de novo* assembly was performed using all non-host data from all root metagenomes. This assembly yielded 1,760,490 contigs larger than 500bp, and this assembly had an N50 value greater than 1,600 bases. In total, there were 6,422,376 predicted genes and a non-redundant gene catalogue (based on 95% similarity) was established with 5,359,885 genes. These genes were mapped to the SEED (28) and KEGG (Kyoto Encyclopedia of Genes and Genomes) (29) databases for functional predictions, and approximately 22% of the non-redundant genes were annotated by each database. For shotgun metatranscriptome analyses, 27-33 million sequences (paired-end) were generated per sample (**Table S1**). Metatranscriptome reads were mapped to the gene catalogue established from the metagenomics analysis (microbial transcriptome, with 11-51% of the sequences mapped) or to the available plant genome (host transcriptome analysis). As the lettuce genome is not fully annotated, sequence data generated from lettuce plants were screened for orthologs of known tomato genes, when possible.

### Host functions actively associated with TWW irrigation

Plant host physiological response to irrigation with water of lower quality has been previously reported, with significantly reduced yield of both plants under TWW irrigation (*e.g.*, (26) **Table S2**). In this study, we performed deep sequencing of plant transcripts to identify stresses that TWW irrigation imposes on plant roots. Across two growing seasons, a total of 45 tomato genes and 645 predicted lettuce transcripts were significantly differentially expressed between irrigation treatments. Of the 645 lettuce transcripts, only 141 could be annotated by comparison with known tomato genes (**Table S3**).

To identify the most robust effects of TWW treatment, tomato and lettuce differentially abundant transcripts were analyzed together. A network analysis of enriched transcripts was performed to predict interactions and highlight clusters of associated genes (**Figure 1**). The FW-enriched gene network consisted of 97 nodes, indicating the number of enriched genes that were identified by the STRING protein-protein interaction network database. Similarly, the TWW-enriched gene network consisted of 86 nodes. The FW-enriched gene network, however, was linked by only seven edges representing predicted protein interactions (direct physical interactions, as well as predicted functional association). In contrast, the TWW-enriched gene network was linked by 69 edges, with a significantly higher number of interactions then expected (p value<0.0001, by Random Graph with Given Degree Sequence (RGGDS)) (**Figure 1a, b**). The TWW gene network of both plants was enriched (Aggregate Fold Change, permutation-based, non-parametric test) primarily with various heat-shock transcripts, including Hsp20, Hsp70, and DnaJ. Heat shock proteins are prevalent in plants and are active during normal growth (30) Such genes also show a stress response, and can also be activated in response to many stress cues, including heat, cold, water stress, salinity, osmotic stress, and oxidative stress (30, 31). In addition, tubulin and ‘FKBP-type peptidyl-prolyl cis-trans isomerase’ genes were also significantly enriched under TWW exposure (**Figure 1b,c**). Tubulin reorganization has been shown under salt stress, cold shock, aluminum exposure, interaction with pathogens and more (32, 33, 34, 35). Overall, the plant response to irrigation with TWW, as detected by transcriptome analysis, was largely restricted to highly general stress response genes that are expressed under a wide range of environmental conditions.

**Figure 1:**
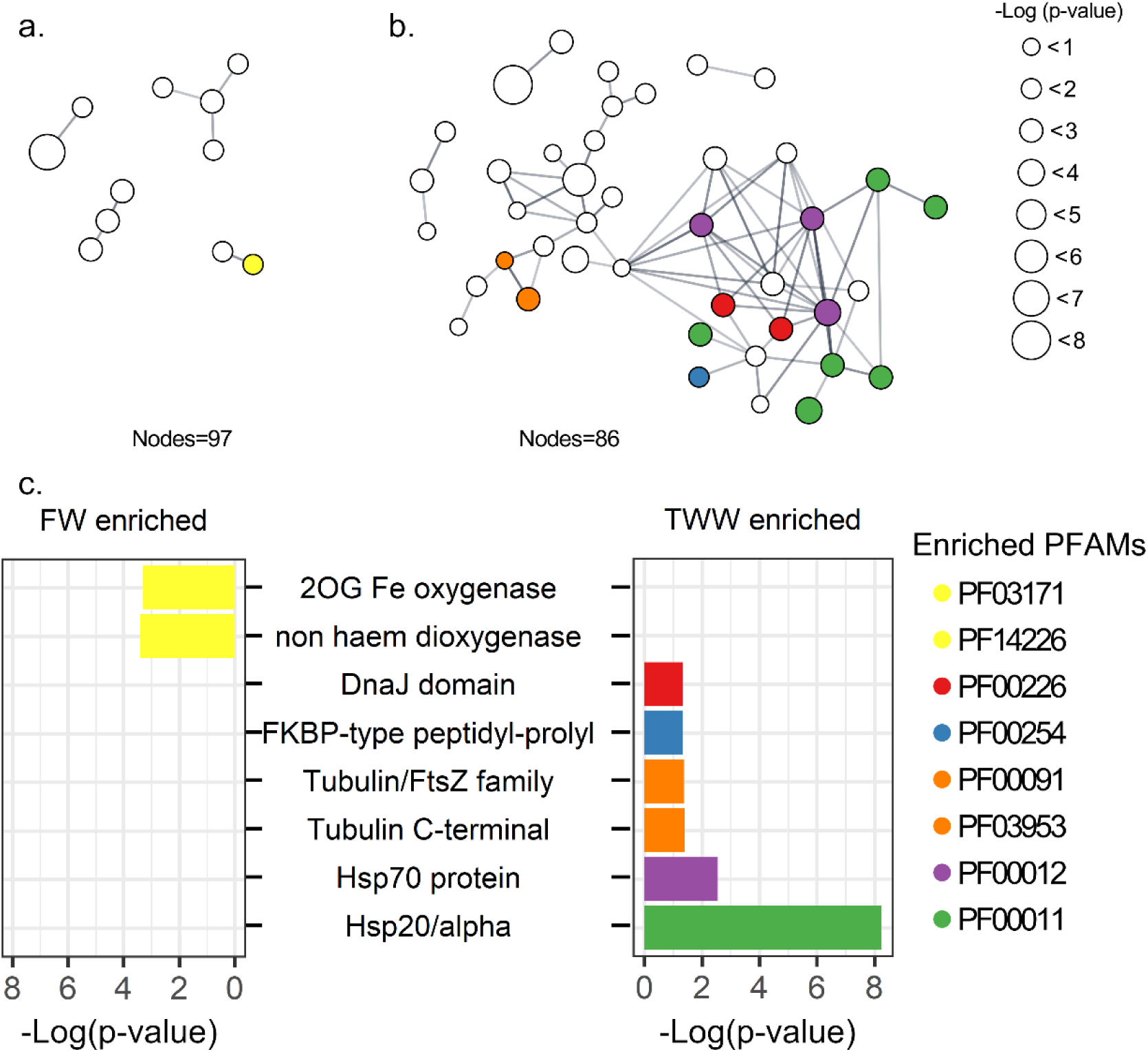
Plant host functional response to TWW irrigation revealed by tomato and lettuce transcriptome analysis. Differentially expressed plant genes between FW or TWW irrigation were determined using the software package EdgeR, with significance set at FDR P<0.05. Lettuce transcripts were annotated by comparison to the tomato genome. STRING protein interaction networks are presented for (a) FW and (b) TWW irrigation enriched gene list. Only nodes linked by edges are shown. The size of each node is proportional to the log10 (p-value) of the enriched gene. Colored are PFAM protein domains significantly enriched within the network landscape. The log10 (p value) of the enriched PFAM domains are presented in (c) FW and (d) TWW enrichment analyses.

### Shifts in microbiome associated with TWW irrigation (DNA-based metagenomics)

#### Functional profiling demonstrates the extent to which root microbiomes respond to environmental factors

The taxonomic affiliation of root-associated microbial communities was determined by analysis of annotated genes from the metagenomes (**Fig. 2f; Table S4, S5**). The vast majority of annotated genes were derived from bacteria (96.7% of all mapped reads), while the percentage of reads derived from Fungi (1.2%), Archaea (0.5%), and viruses (0.13%) was much lower. Despite a prior mapping step to remove host reads, 1.2% of annotated gene counts could still be mapped to plant genomes. The root microbiome was primarily composed of bacteria from the phyla Proteobacteria (44% of all mapped reads) and Actinobacteria (33%).

**Figure 2:**
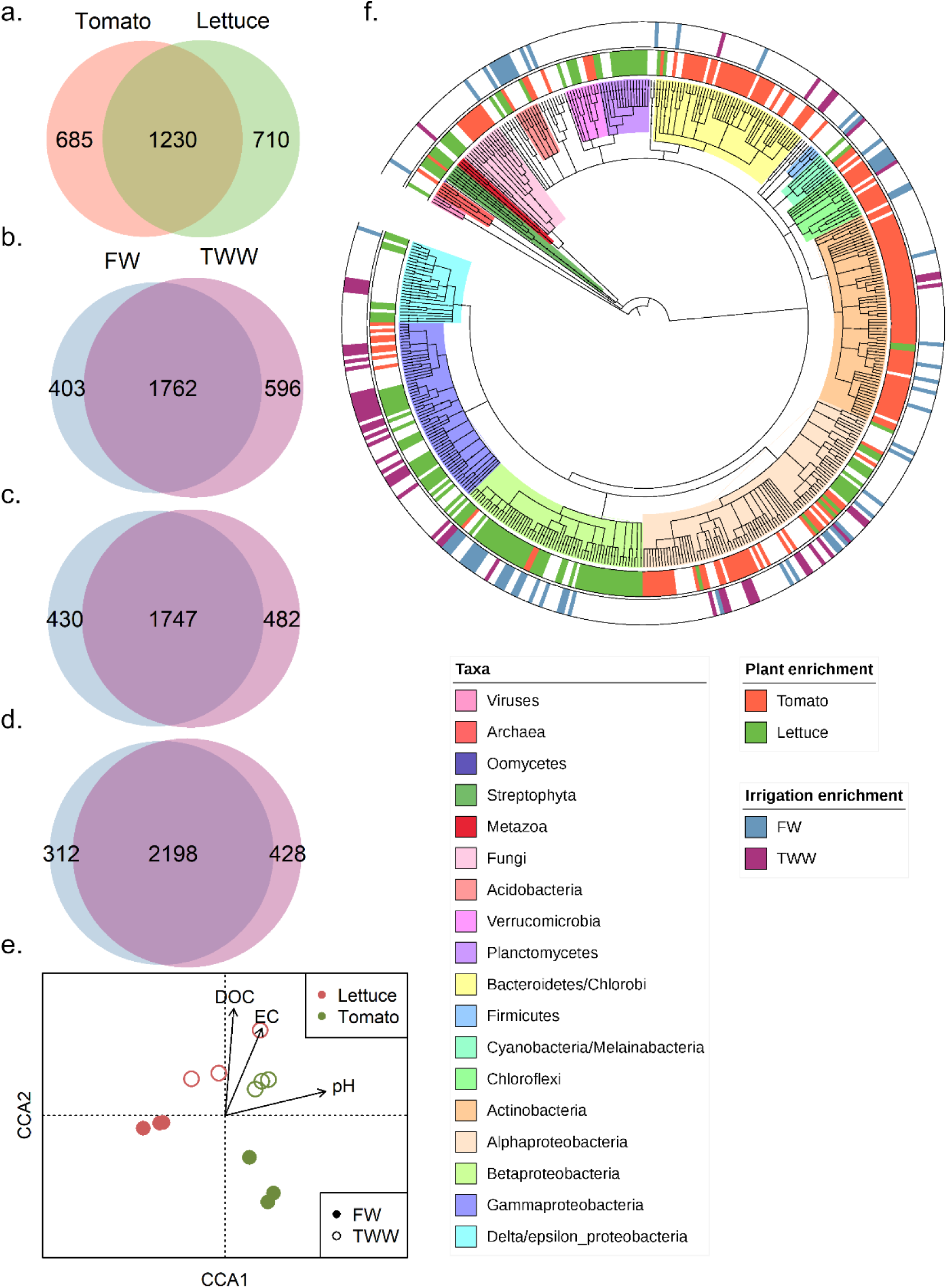
Effect of plant host type and irrigation treatment on root-associated metagenome. (a) VENN diagram of differentially-abundant genes between FW-irrigated tomato and lettuce (DESeq2 FDR P<0.05). VENN diagrams of differentially-abundant genes by irrigation type in (b) tomato or (c) lettuce, and (d) in both hosts combined. (e) Conical correspondence analysis (CCA) of all SEED-annotated genes with DOC, EC and pH as constrained variables. (f) Microbial composition predicted by a least common ancestor (LCA) pipeline (MEGAN6) of the predicted gene catalogue. Taxonomic groups are displayed in the inner ring, and differentially-abundant taxonomic groups between the two tested plant types are highlighted in the middle ring. The outer ring highlights the taxonomic groups that are significantly differentially abundant between irrigation treatments.

The relative abundance of taxa was compared across experimental conditions of plant host type (tomato *vs.* lettuce) and irrigation water quality (FW *vs.* TWW) using DESeq2 method for comparing differential abundant count data (36)(**Figure 2f**). Broadly, 34% of all taxonomic groups (with highest available taxonomic resolution, based on MEGAN6 least common ancestor) were significantly (Wald test, FDR corrected p value<0.05) more abundant in tomato roots, as compared to 31%, significantly associated with lettuce roots. Many taxa from the phlya Actinobacteria and Bacteroidetes/Chloroflexi were significantly more abundant in tomato roots relative to lettuce, while Betaproteobacteria and Planctomycetes were strongly and significantly associated with lettuce roots. Irrigation water quality mostly affected Proteobacterial taxa (10% of microbial taxa were significantly more abundant in TWW-irrigated roots, as compared to 11% of microbial taxa enriched in FW-irrigated roots. 60% of all significantly abundant taxonomic group were identified as Proteobacteria). Acidobacteria and Betaproteobacteria were significantly more abundant in FW-irrigated roots and Gammaproteobacteria significantly more abundant in TWW-irrigated roots (all data available at supplementary **Table S4**).

Root microbial metagenomes from lettuce and tomato were annotated and mapped to the SEED database to identify functional genes significantly associated with plant host type (tomato and lettuce) and irrigation water quality (FW and TWW). A comparison of differentially abundant functional genes between tomato and lettuce root microbiomes, demonstrated strong host specificity in microbiome gene content (**Figure 2a**), consistent with our prior analyses of root microbiomes of different plant species grown in identical soils (19). In this study, greater than 50% of SEED annotated genes (from a total of 2625 “functional role”(28)) were significantly (based on Wald test, adjusted p-value<0.05) more abundant in either tomato or lettuce roots (26% in tomato relative to lettuce and 27% in lettuce relative to tomato; **Figure 2a**).

Irrigation water quality also affected root-associated microbiome functional profile (**Figure 2b, c, d**). Initially, the effect of irrigation type was examined in tomato and lettuce systems independently by comparing SEED-annotated gene abundance using DESeq2 method. Irrigation type determined 36% of the tomato root metagenome (15% of all annotated gene list in tomatoes were significantly more abundant in FW irrigated plants compare to 21% more abundant in TWW irrigated plants, Wald test, **Figure 2b**) similarly to 34% of the lettuce root metagenome (FW- 16%, TWW- 18%, **Figure. 2c**). To identify commonalities in response to irrigation, irrigation effects were examined in a dataset of both plant hosts combined. This combined analysis identified microbiome genes that were positively associated with FW irrigation (11% of all SEED annotated genes) or with TWW irrigation (15% of all SEED annotated genes) (**Figure 2d**). Overall, the combined root microbiome functional profiles were significantly associated with plant host (tomato *vs*. lettuce, F=10.1, p value=0.001) and irrigation treatment (FW *vs*. TWW, F=6.6, p value=0.004) as determined by a PERMANOVA test based on the Bray-Curtis dissimilarity index of the SEED annotated gene counts (**Figure S1**). For further analyses, unless otherwise indicated, ecosystem comparisons of plants grown in FW and TWW were performed on data combined from both plant hosts.

We previously measured significant increases in pH, dissolved organic matter (DOC) and electrical conductivity (EC) in TWW-irrigated soils relative to FW-irrigated soils (26)(also at **Table S6**). Similar patterns were observed in this study through canonical correspondence analysis (CCA) (**Figure 2e**). The CCA presents the relationship of the measured soil parameters and root microbiome functional gene profile (all SEED-annotated gene counts). An ANOVA permutation test was significant (F= 3.2, p value=0.001 for the full model), and the constrained variables (i.e., pH, DOC, EC) accounted for 54.8% of the variance. DOC (F=2.4, p value=0.038) and pH (F=5.7, p value=0.002) were found to significantly explain portions of the variance associated with the observed microbiome functional profile, while EC was not significant (F=1.57, p value=0.163). The microbial community functional gene profiles of FW- and TWW-irrigated root samples were separated along the CCA2 axis, with DOC loading primarily on the CCA2 axis. Conversely, the microbial community functional gene profiles of samples from different plant hosts were separated primarily along CCA1 axis, with pH loading primarily on the CCA1 axis.

#### SEED & KEGG functional categories enriched or depleted in metagenomes of TWW-irrigated roots relative to FW-irrigated roots

SEED- annotated genes highly significantly (p<0.01) associated with irrigation water quality across both plants were examined, and in total 438 genes were identified (**Figure. 3**). Of these genes, 286 were enriched in TWW-irrigated roots and 152 enriched in FW-irrigated roots. These genes were clustered into general categories (SEED level 1, based on the hierarchical clustering available on MEGAN6): *e.g.,* carbohydrates, amino acid derivatives, membrane transports, respiration and regulation of cell signaling (**Figure 3a**). Rare categories (represented by fewer than 5 genes) were removed from the analysis. To compare category enrichment, we examined the proportion of genes enriched for each category (**Figure 3b**). For some gene categories (*e.g.,* cell division, cell cycle or carbohydrates), a similar number of genes were enriched in both TWW- or FW-irrigated roots (*i.e.*, no specific effect of irrigation treatment), while others were more strongly skewed to either FW or TWW. For example, membrane transport and transposable element genes were substantially enriched in TWW-irrigated roots. Conversely, the gene categories of nucleosides and nucleotides and sulfur metabolism, were substantially enriched in FW-irrigated roots.

**Figure 3:**
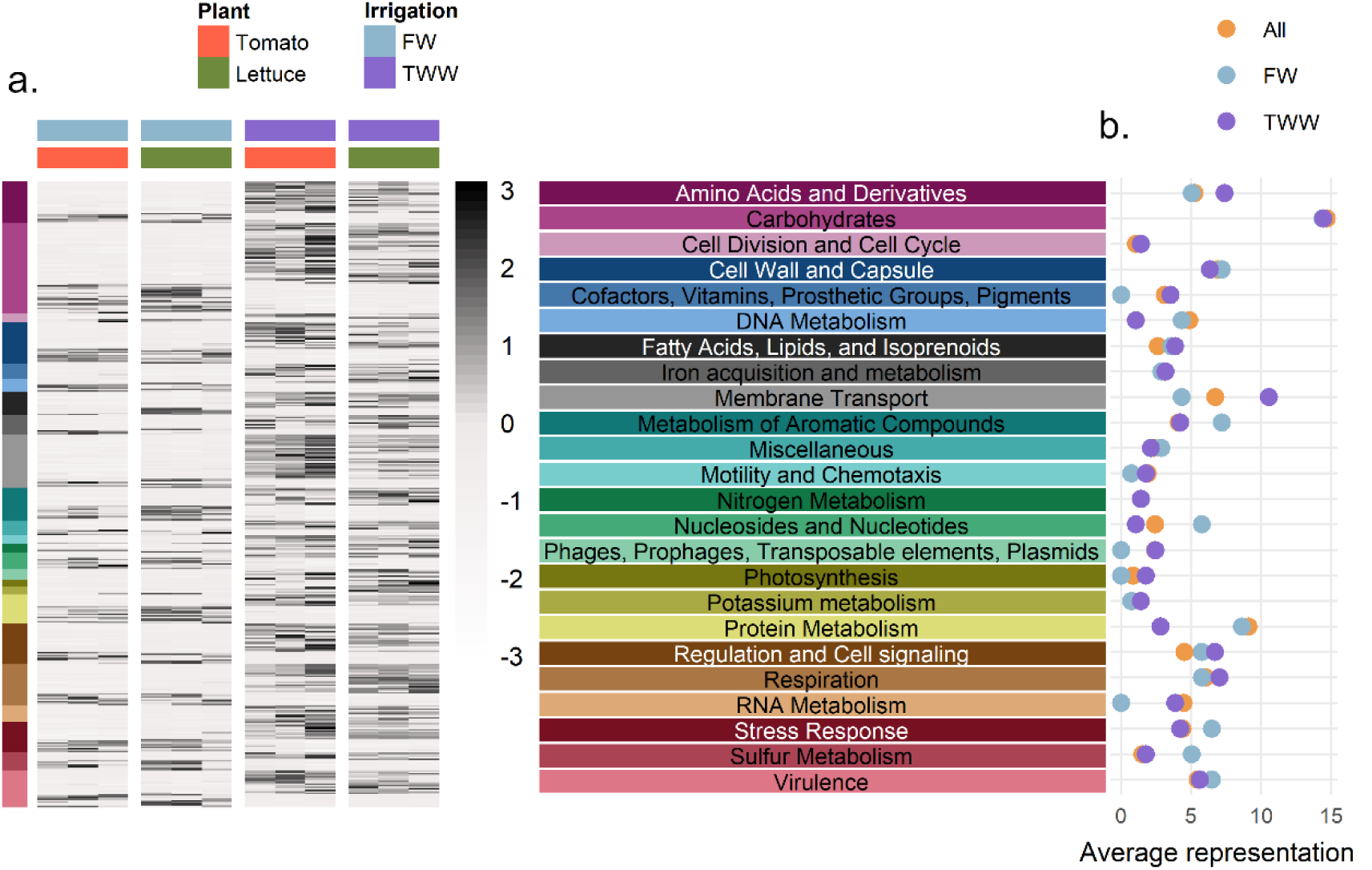
Genetic profile of the 438 significantly differentially abundant genes between irrigation treatments. (a) Heatmap of genes enriched or depleted (FDR P<0.01) in the metagenomes of TWW-irrigated roots (displaying trimmed mean of M values- TMM). Gene abundance was normalized by scaling each row separately. The gene list was clustered to high hierarchy SEED categories. (b) The proportion of genes enriched or depleted in TWW-irrigated root metagenomes compared to the total abundance of that category in all metagenome analysis. Enriched category (TWW, magenta), deprived (FW, light blue), or the proportion within the full gene catalogue (marked as “All”, colored by orange), are highlighted. The proportion was calculated based on the number of gene assigned to the different categories with-in each data set.

An enrichment analysis was also conducted for gene subsystem enrichment and depletion by irrigation method (level 2, based on the SEED hierarchical clustering, tested using Wallenius non-central hypergeometric distribution) (**Figure 4a, Table S7**). One of the most strongly enriched categories (log_2_FC=1.4; relative abundance=0.07%, p value<0.0001) was the Na(+)-translocating NADH-quinone reductase (NQR), a membrane complex that utilizes the respiratory chain to generate a sodium gradient in place of a proton gradient in high pH and sodium conditions (37). In addition, enrichment of multiple membrane-associated subsystems in TWW-irrigated roots was observed, including: (i) sodium-hydrogen antiporter, a common membrane transporter that supports sodium balance in exchange for proton motive force (log_2_FC=1.67; p value=0.002), (ii) pH adaptation potassium efflux system (log_2_FC=1.87; p value=0.007), (iii) mannose-sensitive hemagglutinin, type 4 pilus (MSHA4) (log_2_FC=1.03; p value=0.0009), (iv) alginate metabolism membrane complex (log_2_FC=0.23; p value=0.01). In addition to membrane-associated subsystems, other subsystems were enriched in TWW-irrigated roots, including arginine degradation (log_2_FC=0.56; p value=0.04). Five genes enriched within this subsystem catalyze the complete arginine to glutamate pathway (**Table S7**). Soil Na^+^ concentration, K^+^ concentration and pH were correlated with observed gene abundance patterns (**Figure S2, Table S8**). The relative abundance of TRAP transporters (Pearson’s R_DOM_=0.81, P-value_DOM_=0.001; R_Na_=0.78, P_Na_=0.003; R_K+_=0.81, P_K+_=0.001) and sodium hydrogen antiporters (R_DOM_=0.86, P_DOM_=0.0003; R_Na_=0.78, P_Na_=0.003; R_K+_=0.8, P_K+_=0.001) correlated to organic matter and Na^+^/ K^+^ concentrations. The relative abundance of potassium antiporter genes was correlated with Na^+^ and K^+^ concentrations (R_Na_=0.9, P_Na_<0.0001; R_K+_=0.86, P_K+_=0.0003), and MSHA4 gene relative abundance was correlated with pH (R_pH_=0.88, P_pH_=0.0001). NQR gene abundance was significantly correlated with salt concentration (R_Na_=0.8, P_Na_=0.002; R_K+_=0.81, P_K+_=0.001) and pH (R_pH_=0.79, P_pH_=0.002).

**Figure 4:**
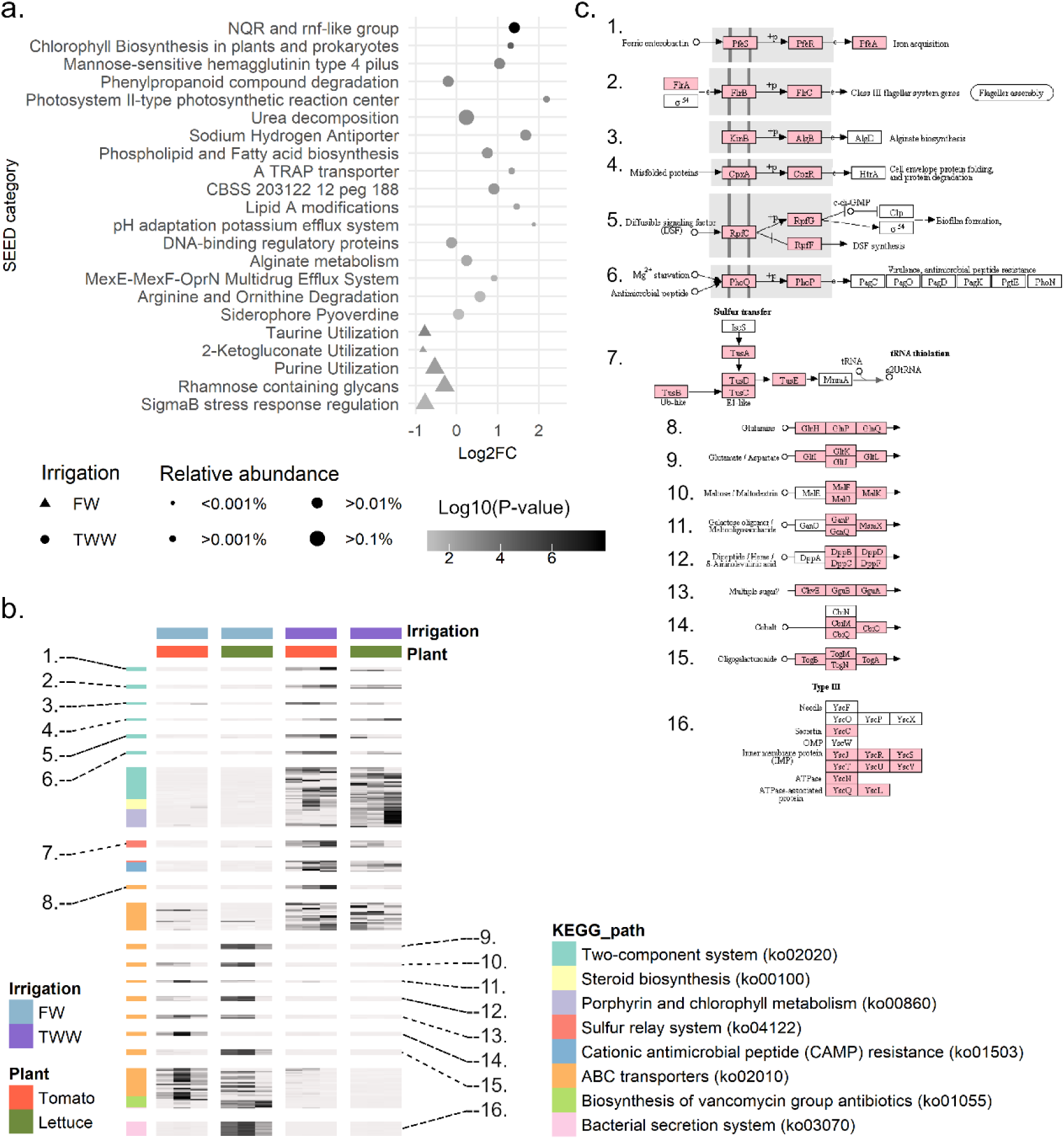
Analyses of SEED subsystems and KEGG pathway significantly enriched or depleted in metagenome of TWW irrigated roots. (a) Dot plot of log2(fold-change) relative abundance of enriched or depleted SEED subsystem. Significantly (FDR P<<0.05, represented by more than two gene families) enriched or depleted gene abundance was computed using the goseq software package, with correction for read counts. Symbols are proportional to the sub-system relative abundance and colored based on the enrichment/depletion log10(p-value). Circles indicate TWW-enrichment while triangles indicated TWW-depleted categories. (b) Heatmap of TWW-enriched or depleted (p-value < 0.05) KEGG pathways (characterized by keggProfiler). Genes of interest, significantly enriched or depleted in TWW-irrigated root metagenomes, are highlighted and colored in pink (c).

Analysis of enriched KEGG pathways (**Figure 4b, c, Table S9**) and modules (**Figure S3**) revealed additional biological processes enriched in TWW-irrigated root microbiomes relative to FW-irrigated root microbiomes. Two-component systems were significantly enriched (40 KEGG genes were enriched, p value<0.0001) in TWW-irrigated root microbiomes relative to FW-irrigated root microbiomes, including pathways involved in misfolded proteins, flagellar assembly, iron acquisition, and Mg^2+^ starvation (**Figure 4c**). In addition, the denitrification gene module was significantly (4 genes, p value=0.006) enriched in TWW irrigated roots (**Figure S3**). Conversely, ABC transporter gene pathways associated with sugar (maltose, galactose and oligogalacturonide), peptide (dipeptide and glutamate/ aspartate) and nutrient (cobalt or nickel) transport were enriched (43 genes, p value<0.0001) in FW-irrigated root microbiomes relative to TWW-irrigated roots. A Type 3 secretion system (T3SS) gene module was also enriched (12 genes, p value<0.0001) in FW-irrigated root communities.

#### Meta-analysis of selected gene counts relative to environmental variables from publicly available metagenomes

A meta-analysis was conducted to establish a global link between metagenome functional gene content and measured environmental variables. We focused on subset of prominent genes from this study that were strongly positively correlated with pH (NQR, Na^+^-H^+^ antiporter) or negatively correlated with oxygen levels (periplasmic nitrate reductase, *napAB*, nitric oxide reductase- *norBC*, nitrous oxide reductase Z, *nosZ*). Metagenomes available at the Joint Genome Institute’s (JGI) Integrated Microbial Genomes and Microbiomes repository (n=14,596) were screened. Environmental pH measurements were available for a subset of these metagenomes (n=1,588), and of this subset, 160 metagenomes had a total number of predicted genes greater than 100,000 (**Table S10**). Within these 160 metagenomes, the relative abundance of genes annotated as NQR (pairwise Wilcoxon rank test, Bonfferoni correction- p value_pH<7: pH7-8_=0.02; p value_pH7- 8:pH>8_<0.0001; p value_pH<7:pH>8_=0.002) and Na^+^-H^+^ antiporter (p value_pH<7: pH7-8_<0.0001; p value_pH<7:pH>8_<0.0001) were strongly correlated with measured pH values (**Figure 5a, b**). Using the same filtering criteria, 257 metagenomes were identified with oxygen measurements and greater than 100,000 predicted genes. In these metagenomes, the abundance of *napAB*, *norBC* and *nosZ*, in the denitrification pathway, were enriched in samples with lower measured oxygen (**Figure 5d, e, f, g**). No such trend was observed for housekeeping genes such as gyrase B (*gyrB*, **Figure 5h**). Salinity or Na^+^ concentrations were measured only in small subset of available metagenomes and were not analyzed further.

**Figure 5:**
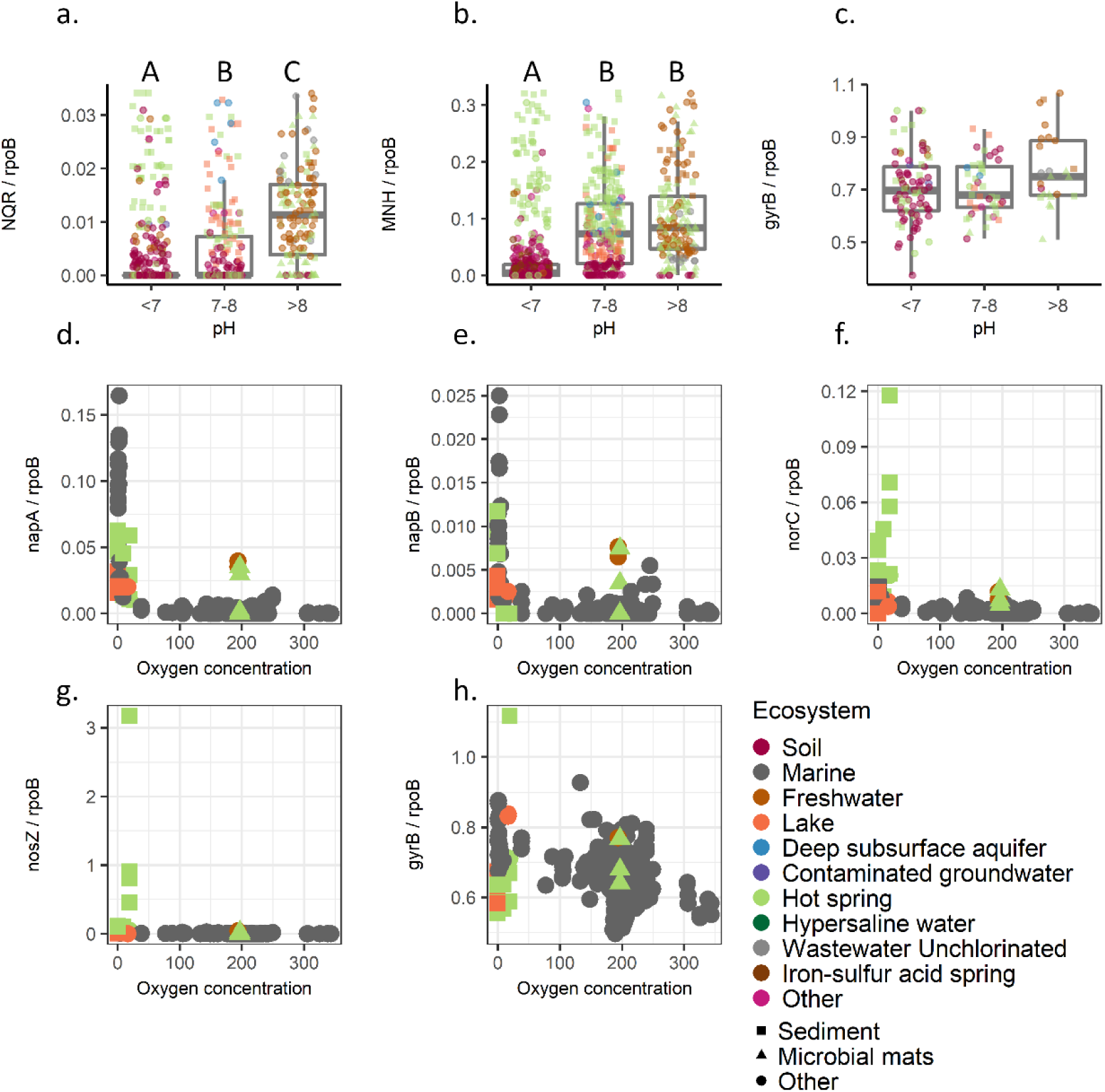
Meta- analysis of selected pH and oxygen responsive genes. Publicly available (by JGI) subset of 160 environmental metagenomes with pH measurements, were screened. Box plot of gene counts in acidic pH (<7), neutral (7–8), or alkaline (>8) pH for (a) NQR operon (6 subunits), (b) Na- H antiporter operon (7 subunits), and (c) *gyrB* as control. Detailed pattern for each subunit is available in Figure S6, S7. Outliers are not displayed (1.5x 0.25-0.75 quantiles). Significant differences in gene counts, by Wilcoxon rank sum test (Bonfferoni correction, P<0.05), are marked in letter report (A, B and C). Oxygen measurements were available for subset of 257 environmental metagenomes. In these metagenomes, the abundance of *napA* (d), *napB* (e) *norBC* (f) and *nosZ* (g) was compared to oxygen levels. (h) *gyrB* was used as control. All gene counts are in proportion to *rpoB* gene.

### Microbial gene expression patterns associated with TWW irrigation (RNA-based metatranscriptomics)

An enrichment analysis of the root-associated microbial metatranscriptomes was performed to identify SEED-annotated genes and subsystems that were significantly differentially transcribed between plants irrigated with TWW relative to those irrigated with FW (**Figure 6, Table S11**). In total, 10.1% of SEED-annotated genes were significantly differentially expressed in roots of TWW-irrigated plants relative to FW-irrigated plants (Wald test, FDR corrected p value<0.05). Specifically, 7.2% of such genes had higher expression and 2.9% lower expression in TWW-irrigated roots relative to FW-irrigated roots. SEED-annotated genes were clustered into 761 SEED subsystems (level 2, based on SEED hierarchical clustering), and of these, 8 were over-represented in TWW-irrigated root microbial communities while only a single subsystem was significantly over-represented in the FW-irrigated root transcriptomes. The most highly and significantly over expressed gene sub-systems in TWW-irrigated roots were NQR, TRAP transporters, sodium-hydrogen antiporters, alginate metabolism genes and MSHA4 (**Figure 6a**). All of these genes were also significantly enriched in metagenomic analysis of TWW-irrigated roots relative to FW-irrigated roots. Genes involved in alginate metabolism were only slightly enriched in metagenomes of TWW-irrigated roots (log_2_FC=0.23, p value=0.01) but were strongly over-expressed in TWW-irrigated metatranscriptomes relative to FW-irrigated roots (log_2_FC=1.7, p value=0.0004). Conversely, the overall expression level of hydrogenase subsystem genes was significantly higher in FW-irrigated roots relative to TWW-irrigated roots (log_2_FC=1.6, p value>0.0001), though their relative abundance at the DNA level was not substantially affected by irrigation treatment (**Figure 6b**).

**Figure 6:**
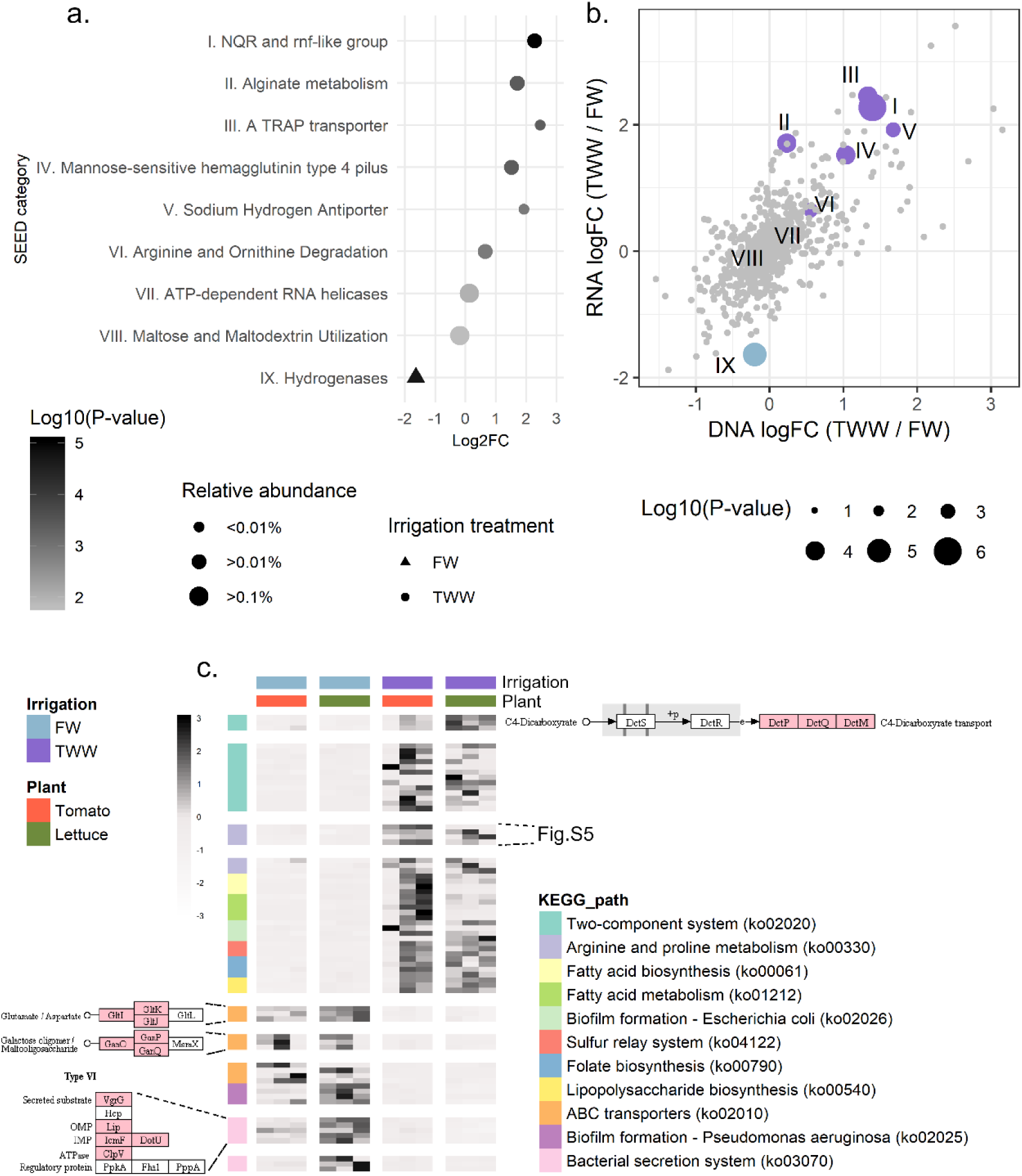
Expressed functions associated with irrigation treatment. (a) Dot plot of Log2(fold-change) SEED subsystems enriched or depleted in TWW irrigated root metatranscriptomes. Significantly (FDR p <0.05, represented by more than two gene families) enriched or depleted transcript abundance was computed using the goseq software package, with corrections for read abundance. Symbols are proportional to the sub-system relative abundance and colored based on the enrichment or depletion log10(p-value). Circles indicate TWW-enriched categories and triangles indicate TWW-depleted categories. (b) Differential metagenome enrichment (TWW/FW fold change) compared to differential metatranscriptome expression level. Highlighted (colored) categories significantly enriched or depleted in the metatranscriptome analysis. ‘ns’= ‘not significant’. Symbols are proportional to the log10(p-value) enrichment in the metatranscriptome analysis. Numbers label the enriched category, as marked in (a). (c) KEGG pathway-level enrichment or depletion in TWW-irrigation root metatranscriptomes (p-value < 0.05), based on keggProfiler enrichment analysis. Significantly enriched or depleted gene clusters in TWW-irrigated roots are highlighted and colored in pink.

Enrichment analysis of KEGG pathways (**Figure 6c**) and modules (**Figure S4**) was performed (**Table S12**). The transcriptome of the microbial community of TWW-irrigated roots was significantly enriched in 2.75% of all KEGG-annotated genes. In addition, 8 KEGG pathways and 2 KEGG modules were significantly enriched in TWW-irrigated root metatranscriptomes relative to FW-irrigated root metatranscriptomes. In contrast, the transcriptome of the microbial community of FW-irrigated roots was significantly enriched in 1.8% of all KEGG-annotated genes. In addition, 3 KEGG pathways and 3 KEGG modules were significantly enriched in FW-irrigated root metatranscriptomes relative to TWW-irrigated root metatranscriptomes. This analysis revealed higher microbial relative expression of two-component systems, including the C4 dicarboxylate gene cluster, in TWW-irrigated roots relative to FW-irrigated roots (**Figure 6c**). Moreover, in TWW-irrigated roots, higher relative expression of arginine and proline metabolism genes, particularly those in the arginine-to-spermidine pathway, was observed (**Figure S5**). The relative expression level of ABC transporter genes, including glutamate and galactose transporters, were higher in the metatranscriptomes of FW-irrigated roots relative to TWW-irrigated roots (10 enriched genes, p value=0.0004). The abundance of type 3 secretion system (T3SS) genes was significantly higher in the metagenomes of TWW-irrigated root microbial communities relative to those of the FW-irrigated root metagenomes. However, the level of expression of type 6 secretion system (T6SS) genes was most highly expressed under FW-irrigation conditions (7 enriched genes, p value<0.0001).

## Discussion

We previously studied the effect of TWW irrigation on soil and root microbial community structure and composition (26). In that study, irrigation water quality and soil type were major explanatory variables for the observed soil microbial community structure and were of a similar magnitude. Similarly, the effect of irrigation water quality on root microbial community structure was of a similar magnitude to the plant host effect (26), demonstrating the responsiveness of the microbial community to both host and environmental factors. In the current study, we have attempted to harness the rhizoplane microbiome – existing at the interface between the plant and the surrounding soil – as a sensor for detecting *in situ* environmental conditions at the plant-soil interface, including factors leading to host stress. The main incentive in using the host-associated microbiome as a biosensor lies in the fact that high resolution is desired for accurate definition of the factors contributing to host physiological status (38). Comparing the differences in the relative abundance of microbial genetic features (*i.e.,* metagenome analysis) or expression of microbiome genes (*i.e.,* metatranscriptome analysis) can aid in the identification of long-term stressors imposed on the host under these conditions as well as short-term stressors revealed by expression of genes processing environmental cues at time of sampling. Analyses can be performed at different levels of hierarchical gene annotation and can be performed using gene level annotation (*e.g.,* SEED database) and enriched pathways or modules (*e.g.*, KEGG annotation).

Most commonly used methods for studying root- soil interface employs microelectrodes (39), or specific dyed root imaging in “rhizoboxes” (40, 41). Both methods measure only pre-defined variables, eliminating the possibility of discovering novel or unsuspected stressors. Moreover, experimental design forces manipulating natural environment by growing plants in designed cells or by removing plants from soil for further experimental procedure. Furthermore, in studies where transcriptional response is examined, plant host response is often tested under severe stress in unnatural short term experimental design (42). We sought to be able to assess environmental factors leading to plant physiological status under more natural agricultural conditions.

A secondary motivating factor for the use of the microbiome as a biosensor lies in the observed low-resolution response of the host organism. In this study, upregulation of stress response genes was identified in the transcriptome of host roots irrigated with TWW relative to those roots irrigated with FW. However, the specific nature of the stresses remained unresolved, with transcriptome analysis revealing only the differential expression of genes involved in a highly general stress response associated with heat shock proteins (43, 44). In fields, plants are expose to myriad fluctuating biotic and abiotic environmental conditions, which force plants to tailor their gene transcriptional profiles. Therefore, individual abiotic stress response cannot be extrapolated to plant experiencing multiple stress conditions. The nature of the stress cannot be predicted based on experimental profiling of individual stress response under regulated conditions (45, 46). Moreover, other types of stress regulation can also mediate plant response, including post-transcriptional regulation of RNA by micro RNAs (miRNAs) or other small noncoding RNAs (47), and through protein modification (48). Such regulation is not as easily measured as gene expression.

In contrast to the host, the genetic diversity of the host-associated microbiome is much greater (27), and the extraordinarily high microbial diversity in soils provides the plant with a wide selection of organisms competing for access to root exudates (18). While the plant host can alter gene expression profiles in response to changing environmental conditions, both the membership of the root-associated microbial community and the expression patterns of the root-associated microbial community can be altered. Thus, the microbiome provides us with a highly dynamic and sensitive target with the potential for both short-term responsiveness (*i.e.*, metatranscriptome) and long-term responsiveness (*i.e.*, metagenome). In this study, we observed that in response to long-term irrigation with TWW, both the metagenome and metatranscriptome were significantly altered. Statistical analysis of microbial features lead to the identification of significantly differently abundant genes, gene transcripts and pathways. Some gene of interest, being most significantly enriched, or with known and informative supported data, are presented in **Figure 7**. Critically, the identification of the differentially abundant or expressed microbial features was consistent with the known key stresses imposed by TWW irrigation on the microbial community and the host.

**Figure 7:**
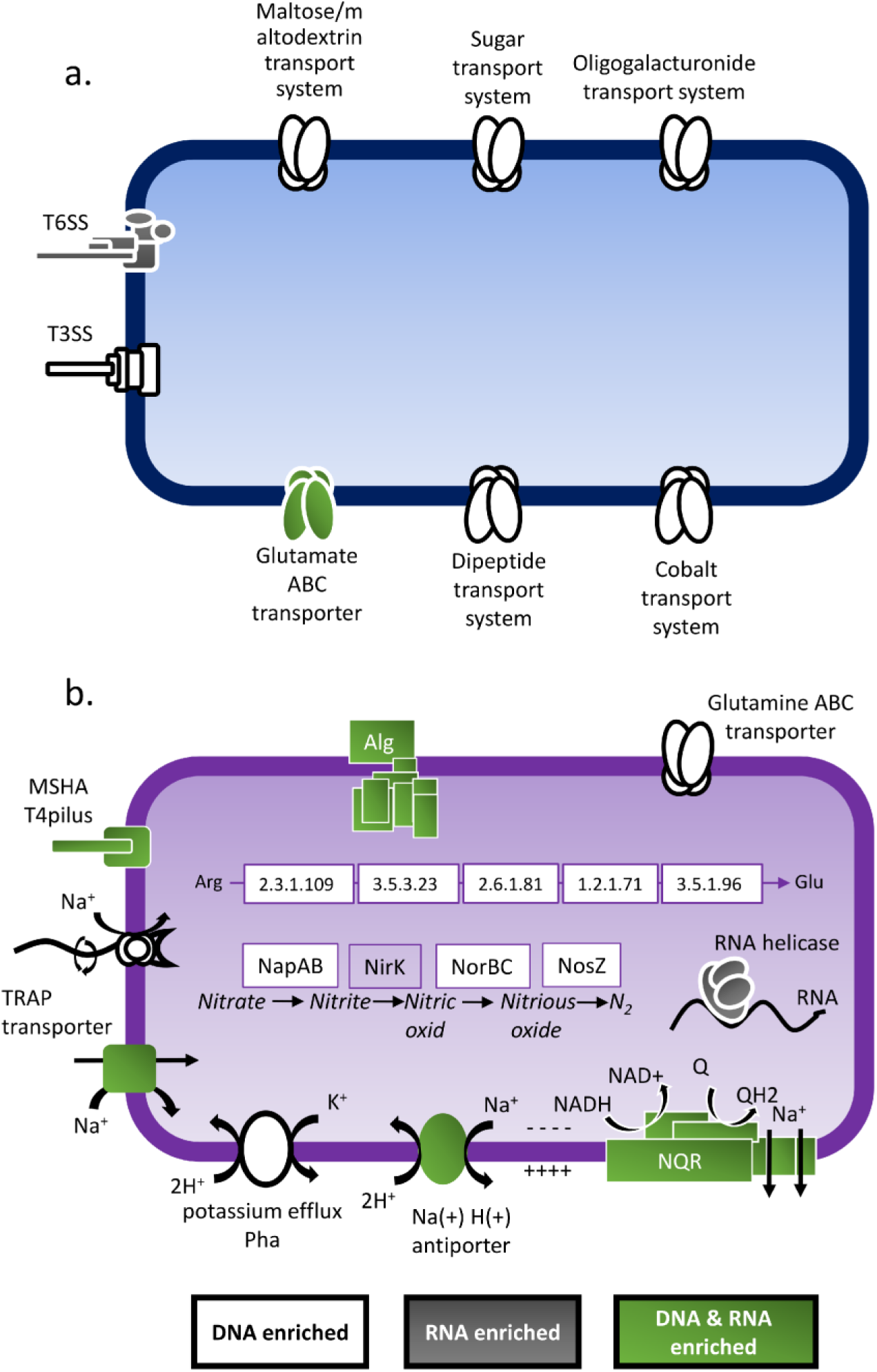
Conceptual model of hypothetical bacteria harboring physiological features (i.e., genes, pathways and modules) enriched in (a) TWW- or (b) FW-irrigated root microbiomes. White symbols indicate features that are significantly enriched at the DNA level (metagenomes), grey features are highly expressed (metatranscriptomes), and green features are significantly abundant and expressed in one treatment relative to the other.

### Level of pH and salinity

Cytoplasmic pH in microorganisms must be at a range suitable for maintaining protein integrity. Bacterial cytoplasmic pH lies at a pH range of 7.4–7.8 (49). In alkaline environments, organisms deploy various mechanisms to maintain intracellular pH and preserve electrochemical gradient in the presence of low proton concentration. To prevent proton loss in alkaline environments, an increase in cytoplasmic pH is achieved by reducing the activity of the proton pumping machinery of the cell respiratory chain. An increase in intercellular proton concentration is attained by an increase in the activity of proton-cation antiporters, reducing the pH gradient along the cell membrane, but increasing the transmembrane electrical potential (50). Under such conditions, some bacteria form a transmembrane sodium gradient, alternatively or concomitantly with a proton gradient. In our plant system, the abundance and expression of Na^+^-transporting NADH:ubiquinone oxidoreductase (*nqr*) genes was significantly enriched under TWW irrigation relative to FW irrigation. In many alkaline environments, NQR constructs the primary sodium efflux system through oxidation of NADH and reduction of quinone. This process creates an electrochemical gradient of net negative charge in the cytosol (51), and the gradient is used by cation/proton antiporters (*e.g.,* Na^+^/H^+^, and K^+^/H^+^) to exchange non-balanced movement of positive charge (H^+^) to the cell (more protons enter the cytoplasm as compared to the efflux of sodium or potassium ions (49, 51, 52, 53). In a global mapping of soil bacterial communities, cation/proton antiporters were observed to be key genes overrepresented in dryland soil, presumably due to the high levels of salt and pH in arid soils (54). Additionally, we measured an increase in the relative abundance and expression of Tripartite ATP-Independent Periplasmic (TRAP) transporters in TWW roots. TRAP transporters have been recently demonstrated to use a membrane-associated sodium gradient to facilitate transport of ligands (55, 56, 57). Similarly, the increase in abundance in flagella assembly genes under TWW irrigation conditions may also suggest the use of sodium motive force for flagella performance (58, 59). This is further consistent with cell motility as a critical feature of rhizosphere competence (19). The signs for high pH stress obtained by the metagenome and metatranscriptome analysis are consistent with soil chemical analysis, as elevated pH conditions can result from long-term TWW irrigation.

Bacterial adaptation to alkaline conditions is frequently dependent on salt concentration, and elevated salt levels are also found in TWW and TWW-irrigated soils (60). Elevated soil salinity can develop through long-term TWW irrigation, and can adversely affect protein and cell membrane stability. Commonly, microorganisms adapt to high levels of salts through osmoregulation by synthesizing organic solutes, thereby avoid salt imbalance and the influx of toxic salts. More rarely, in stable saline environments, some halophiles mitigate salinity levels through adapting the cell enzymatic activity to the high ionic strength. Both strategies require stabling salt concentration in the cell, mostly by regulating cation proton antiporter activity (61). Efflux of sodium ions by NQR activity and the activation of cation/proton transporters demonstrate that the TWW-irrigated root microbiome and the plant roots are indeed exposed to elevated salinity as compared to the FW-irrigated roots. This finding is consistent with the measurement in this study of higher levels of Na^+^ in the leaves of TWW-irrigated tomato and lettuce plants relative to FW-irrigated plants (**Table S2**) and in other plants (62).

To the best of our knowledge, no prior study has linked environmental microbiome functions to pH level or salinity. However, the association between our significantly differentiated genes to processing specific environmental conditions is established *in vitro* in numerous studies (51, 63). Here, we hypothesized that differentially abundant genes could be used as predictive markers of environmental cue, and this hypothesis is supported by meta-analyses demonstrating a link between single isolate studies and microbial communities *in vivo*. A greater effort in collecting, publishing and making metadata more accessible for metagenomics surveys, will better aid the interpretation and modeling of microbiomes processing environmental conditions.

### Oxygen levels

Microbial gene content and expression patterns have great potential for identifying oxygen conditions *in situ*. We previously demonstrated that different plants, even when grown in the same soil, may produce highly different rhizoplane oxygen conditions, and that shotgun metagenome and metatranscriptome analysis revealed differential expression of denitrification genes and catalase genes (19). In this study, we observed an enrichment in denitrification genes in TWW-irrigated root metagenomes relative to FW-irrigated metagenomes, possibly suggesting a lower oxygen concentration under TWW-irrigation (63). However, the expression level of denitrification genes was not significantly higher in TWW-irrigated roots relative to FW-irrigated roots. This difference in enrichment between metagenome and metatranscriptome could be due to enrichment in the TWW-irrigated rhizoplane of facultative denitrifying microorganisms, with rhizosphere selection based on other physiological capabilities (*e.g.*, motility). Conversely, the lack of enrichment in denitrification genes in TWW-irrigated metatranscriptomes could be a result of time of sampling. The root microbiome functional profile is expected to fluctuate by diurnal or hydration-dehydration cycles (64, 65, 66). Therefore, gene abundance is indicative of the chronic, long term exposure to stress imposed by TWW, while expression levels may represent transient conditions. Plants in this study were subject to twice-daily irrigation and irrigation conditions in TWW deliver higher levels of organic matter and this may lead to localized oxygen depletion (67). However, at the time of sampling, oxygen levels may have increased. Further short-term longitudinal analysis will be required to demonstrate diel- and irrigation-derived shifts in denitrification gene expression patterns.

### Bacterial life style

We observed a significant enrichment of genes associated with surface attachment, colonization and biofilm formation in TWW-irrigated root microbiome. These enriched microbial features included genes encoding for flagella and MSHA type 4 pili; both features have been previously demonstrated to facilitate near-surface motility and bacterial attachment (68). Furthermore, an increase in the relative abundance and expression of alginate producing genes, which catalyze the formation of extracellular polysaccharide matrix in biofilms of many bacterial clades (69), was observed. These results further point to the critical importance of both motility and attachment for rhizoplane microorganisms, as has been previously indicated (70). The reason for significant enrichment of these genes in TWW-irrigated roots relative to FW-irrigated roots is not entirely clear but may be due to overall elevated organic matter in TWW (71). While the focus in this study has been largely on genes enriched in TWW-irrigated systems, several key ABC transporters were depleted in the metagenomes of roots of both plant types irrigated with TWW relative to those irrigated with FW. These genes include transporters for oligogalacturonide, maltose, general sugar and glutamate. The change in relative abundance of these transporters may indicate differential pattern of root deposits, as was observed in roots from cucumber and wheat (19). Plant glutamate secretion patterns have been shown to be mediated by external cues such as salinity, oxidative stress and availability of nutrients (72). Glutamate ABC transporters were depleted, while glutamine were enriched in TWW irrigated roots.

### Conclusions

We hypothesized that the microbial functional gene profiles and expression patterns can serve as *in vivo* sensors of environmental factors affecting hosts and host-associated microbial communities. Environmental surveys or host-associated microbiome analyses frequently yield contradictory or context-dependent results, making the predictive power of such observations inconclusive. Studying the microbiome as a functional unit reacting to a specific environment, however, constitutes a non-deterministic approach, thereby eliminating the need for marker features (*e.g.*, genes, pathways or specific taxa) associated with specific conditions. As the host and its microbiome are similarly exposed to environmental conditions, genetic profiling and expression analysis of the microbiome may be used as a predictive tool to identify stresses affecting hosts. In this study, we employ a well-defined plant host-microbiome system under experimental treatment with FW- or TWW-irrigation, but this approach may be used to define microscale conditions in other host systems.

## Materials and methods

### Experimental design; Mesocosm scale experiment

Tomato (*Solanum lycopersicum*-Heinz 4107) and lettuce (*Lactuca sativa*-Romaine-Assaph) were grown in lysimeters (0.5 m3) for 98 and 42 days, respectively, at Kiryat-Gat-Lachish agricultural research station, northern Negev, Israel (31.605760, 34.791179). The lysimeters were filled with loamy sand soil collected from western Negev, Israel (31.351722, 34.403471). Plants were drip irrigated for 8 summers with fresh water (FW) or tertiary treated wastewater (TWW), derived from the Kiryat Gat wastewater treatment plant (WTP). At harvest, roots were collected, vigorously washed, dried, and frozen on site for further procedures. All samples were composites, consisting of 2-4 plants collected from separate lysimeters, except for FW irrigated lettuce plants which were collected from two lysimeters and were composites of 2 plants each. Detailed procedures, including soil and plant measurements, were described previously (26).

### DNA and RNA extraction

Standard phenol-chloroform nucleic acid extraction protocol was employed for DNA and RNA isolation (26, 73). In brief, 0.2 gr of roots were moderately bead beaten for 45 s at low speed (4.5 m/s) by Fast Prep FP120 (Savant Instruments Inc., Holbrook, NY, USA) with phenol, phosphate buffer pH 8 (with additional 10 μl ml−1 β-mercaptoethanol -Sigma-Aldrich, St Louis, MO, USA) and 1.25% CTAB (Hexadecyltrimethylammonium bromide, Sigma Aldrich). Following phenol-chloroform wash, nucleic acids were precipitated with polyethylene glycol (PEG) and ethanol. Nucleic acids were split for DNA and RNA isolation. RNA samples were treated with RQ1- DNase (Promega, Madison, WI, USA) and complete DNA removal from RNA samples was validated by real time reverse transcription PCR. RNA integrity was evaluated with Agilent TapeStation (Santa Clara, CA, US). Ribosomal RNAs were removed using the Ribo-Zero rRNA Removal Kit (Illumina, San Diego, CA), combining bacteria and plant probes. Double- strand complementary DNA (cDNA) synthesis was conducted by Maxima H Minus Reverse Transcriptase (Thermo Fisher Scientific, Waltham, MA USA).

### Library prep and sequencing

Shotgun metagenome libraries were generated using a Nextera XT library preparation kit according to the manufacturer’s instructions (Illumina). Complementary DNA for transcriptome analysis was sheared using a Covaris S2 acoustic device, and libraries were generated using a Accel-NGS 1S Plus DNA Library Kit (Swift Biosciences, Ann Arbor, MI) according to the manufacturer’s instructions. Libraries were pooled sequenced using high-output flow cells with paired-end 2x150 base reads on an Illumina NextSeq500 sequencer. Library preparation and sequencing was performed at the University of Illinois at Chicago Sequencing Core (UICSQC).

### Bioinformatic analysis

Quality control of raw double-strand FASTQ sequences was evaluated by FASTQC software (74), and adjusted by Trimmomatic (75) with customized parameters set to: SLIDINGWINDOW:4:15 MINLEN:100 CROP:145 HEADCROP:15.

### Metagenome analysis

Host sequences were removed by comparing quality checked reads to host genome (tomato or lettuce) with bowtie2 (76), and subsequently removing the reads with SAMtools (77). Metagenomics reads from all three replicates were de novo assembled together with metaSPAdes (78). Gene prediction was performed on scaffolds using the software package Prodigal (79). Predicted genes from all samples were combined, and a non-redundant gene catalog was established based on 95% similarity, using CD-HIT (80). The gene catalog was aligned to the NCBI non-redundant protein database using the software package DIAMOND in sensitive mode (81). Sequence annotation (SEED- (28) and Kyoto Encyclopedia of Genes and Genomes- KEGG- (29)) and predicted taxonomy were achieved with MEGAN V6 (82). To attain count data (number of mapped read for each gene), quality checked reads (after host read removal) were aligned to the annotated gene catalog by bowtie2, while analogous read annotated terms were summed using a custom python script.

### Metatranscriptome analysis

Quality checked RNA reads were aligned to the gene catalog established from the metagenomics analysis, in a similar fashion to metagenomics count data.

### Host RNAseq

An estimation of transcript abundance for tomato root samples was obtained by aligning quality checked sequences (prior to host reads removal) to the predicted *Solanum lycopersicum* transcripts with Trinity RSEM transcript quantification method (83). Lettuce transcripts first predicted by Tophat and cufflinks for transcript prediction (84), than infered to ortholog tomato genes by OrthoFinder. Transcript quantification was done following similar analysis as for tomato samples.

### Statistical analysis

Metagenome and metatranscriptome statistical enriched gene list (SEED or KEGG annotated) or taxonomic groups were obtained by DESeq2 (36) and compared using the VennDiagram R package (85). Taxonomic trees were visualized using the interactive tree of life (86) and applying the least common ancestor MEGAN algorithm. SEED subsystems enrichment analysis was conducted with the ‘R’ goseq package (87), normalizing to SEED counts. KEGG pathway and module enrichment were analyzed by clusterProfiler package in R (88). Statistical test (MANOVA, ANOSIM) were conducted in R package ‘vegan’ (89), and figures were plotted with R ‘ggplot2’ (90) or ‘pheatmap’ (91). Differentially expressed host transcripts were obtained using the EdgeR ‘R’ package (92), followed by annotation and visualization using the STRING network (93, 94) Cytoscape integrated application (95) for both plant hosts combined. The minimum required interaction score was customized to medium confidence (0.4), and PFAM protein domain enrichment was set to a false discovery rate p value of 0.05.

## Acknowledgements

The authors thank Dr. Nesli Tovi and Dr. Sammy Frenk from the Agricultural Research Organization – Volcani Center. The authors thank Dr. Jonathan Friedman from the faculty of agriculture, Hebrew university. The authors would also like to thank Dr. Maya Ofek Lalzar from Haifa University. This research was supported by research grant no. IS- 4662-13 from the Binational Agricultural Research& Development Fund (BARD), research grant no. 821-0142-13 from the Israel Ministry of Agriculture and Rural Development and USAID-MERC research grant no. M34-011.

