## Supplementary figures for "The microbiome as a biosensor: functional profiles elucidate hidden stress in hosts"

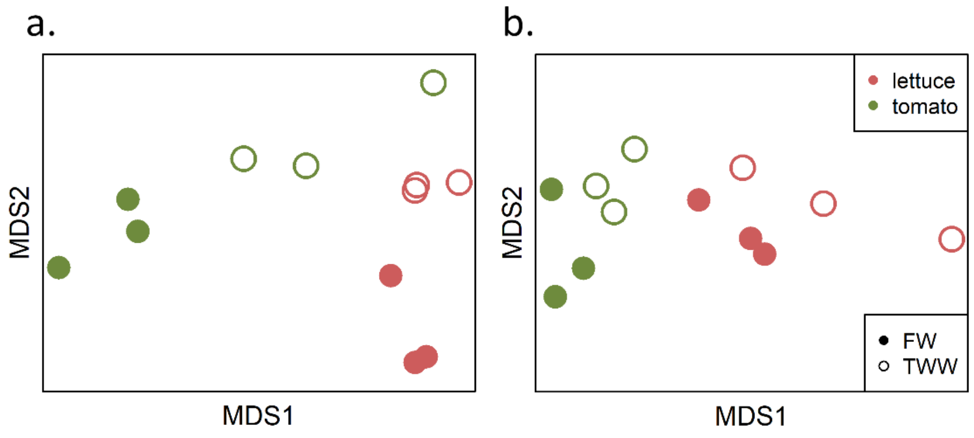


**Figure S1:** Non-metric multidimensional scaling (nMDS) based on the Bray-Curtis dissimilarity index of the SEED annotated gene counts for (a) metagenome, and (b) metatranscriptome.

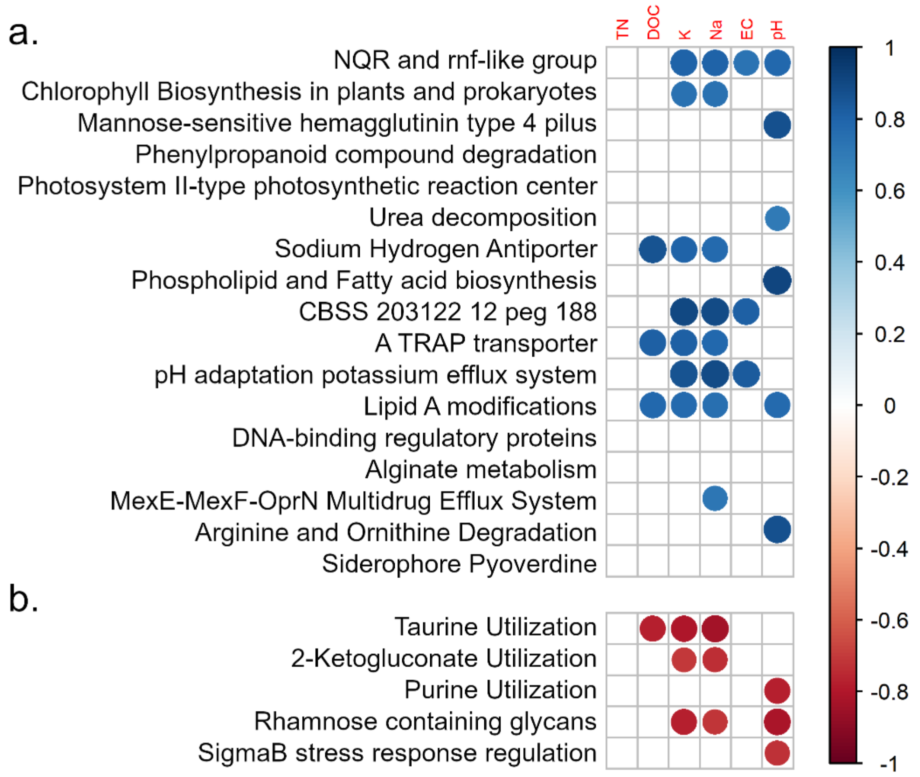


**Figure S2:** Correlation between metagenomics enriched SEED categories and soil environmental parameters, for (a) subsystems enriched in TWW irrigated roots, or (b) enriched in FW irrigated roots.

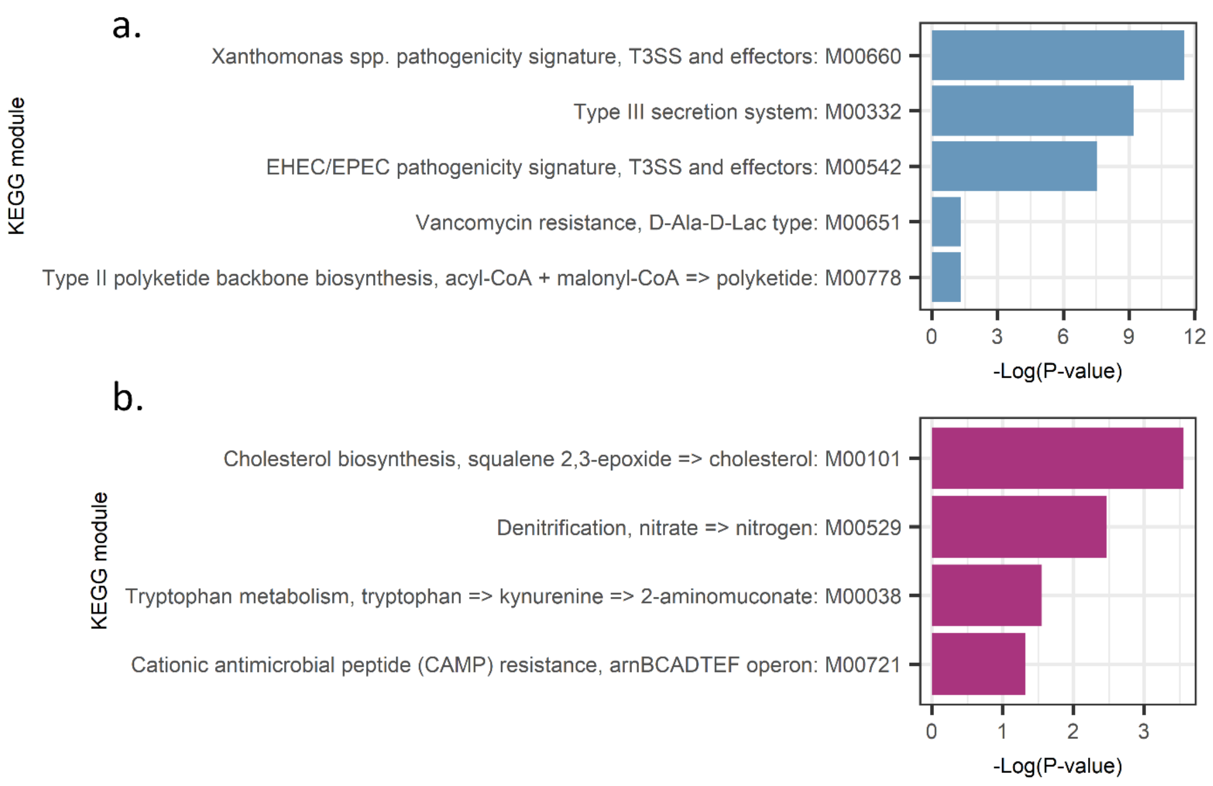


**Figure S3:** Metagenomic enriched KEGG modules in (a) FW irrigated roots, or (b) TWW irrigated roots. The bar chart present the $log_{10}$(P value) of the enrichment analysis calculated by KeggProfiler 'R' package.

a.

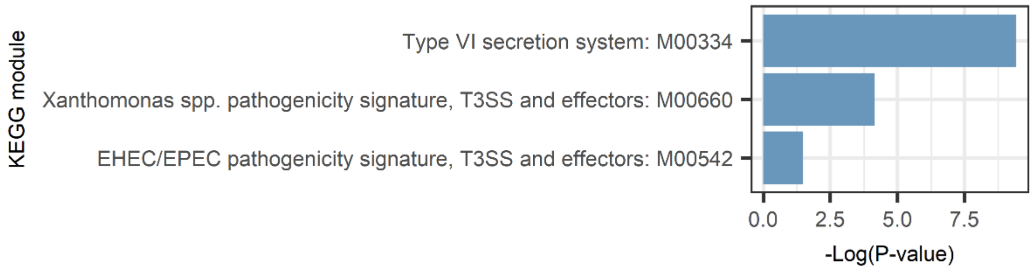


b.

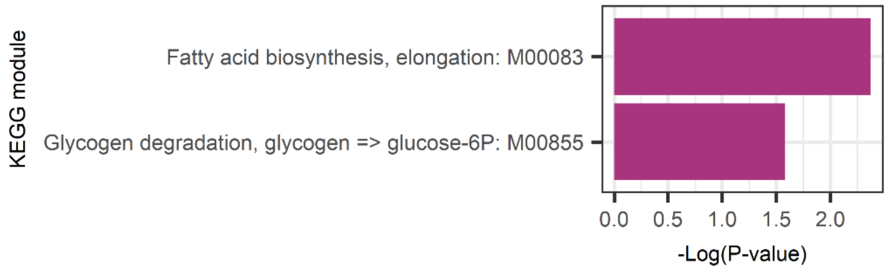


**Figure S4:** Metatransctiptome enrichement KEGG modules in (a) FW irrigated roots, or (b) TWW irrigated roots. The bar chart present the $\log_{10}$(P value) of the enrichment analysis calculated by KeggProfiler ‘R’ package.

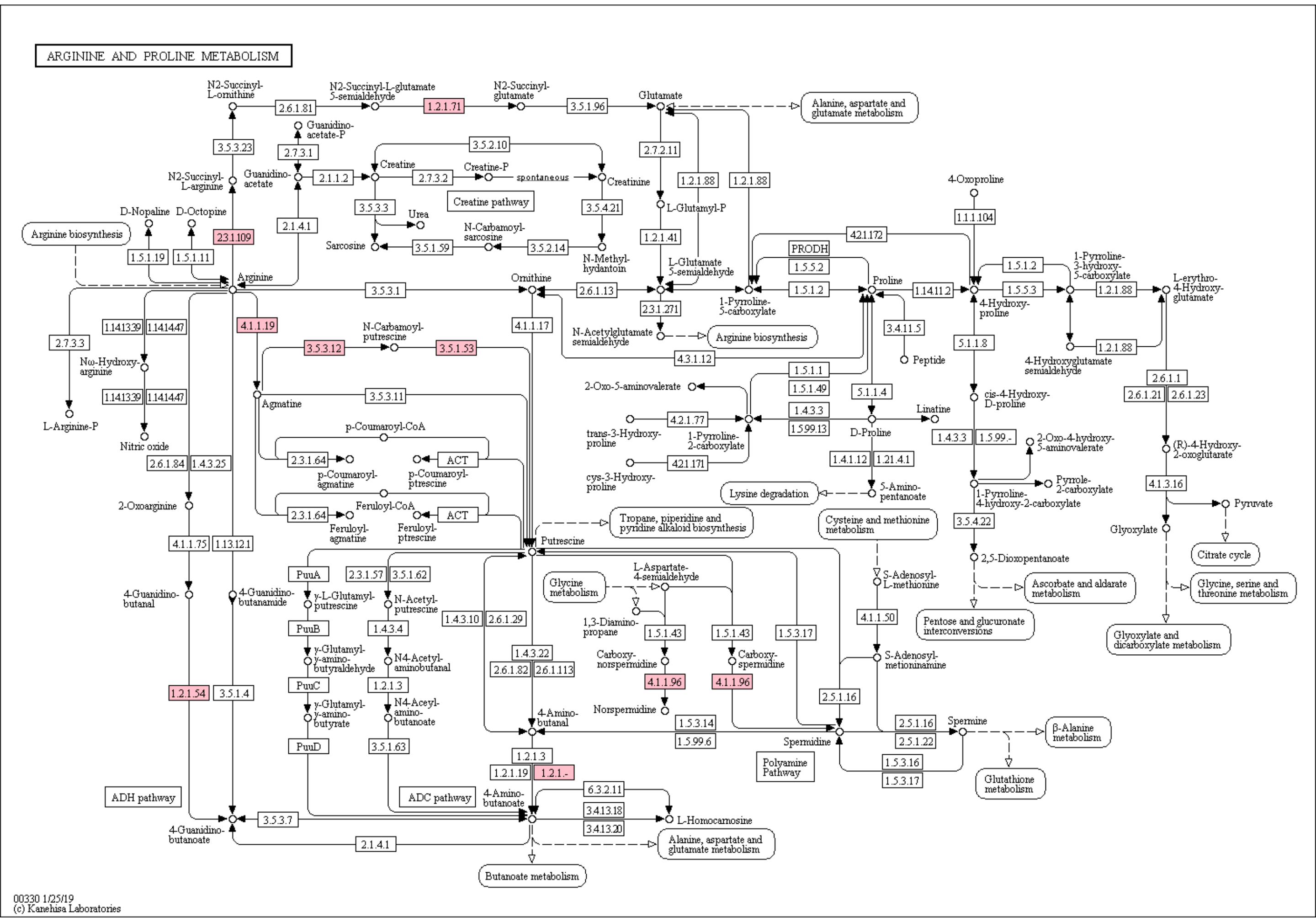


**Figure S5:** Microbial gene expression patterns revealed TWW irrigation enrichement of arginine and proline metabolism genes, particularly those in the arginine-to-spermidine pathway (colored in pink).

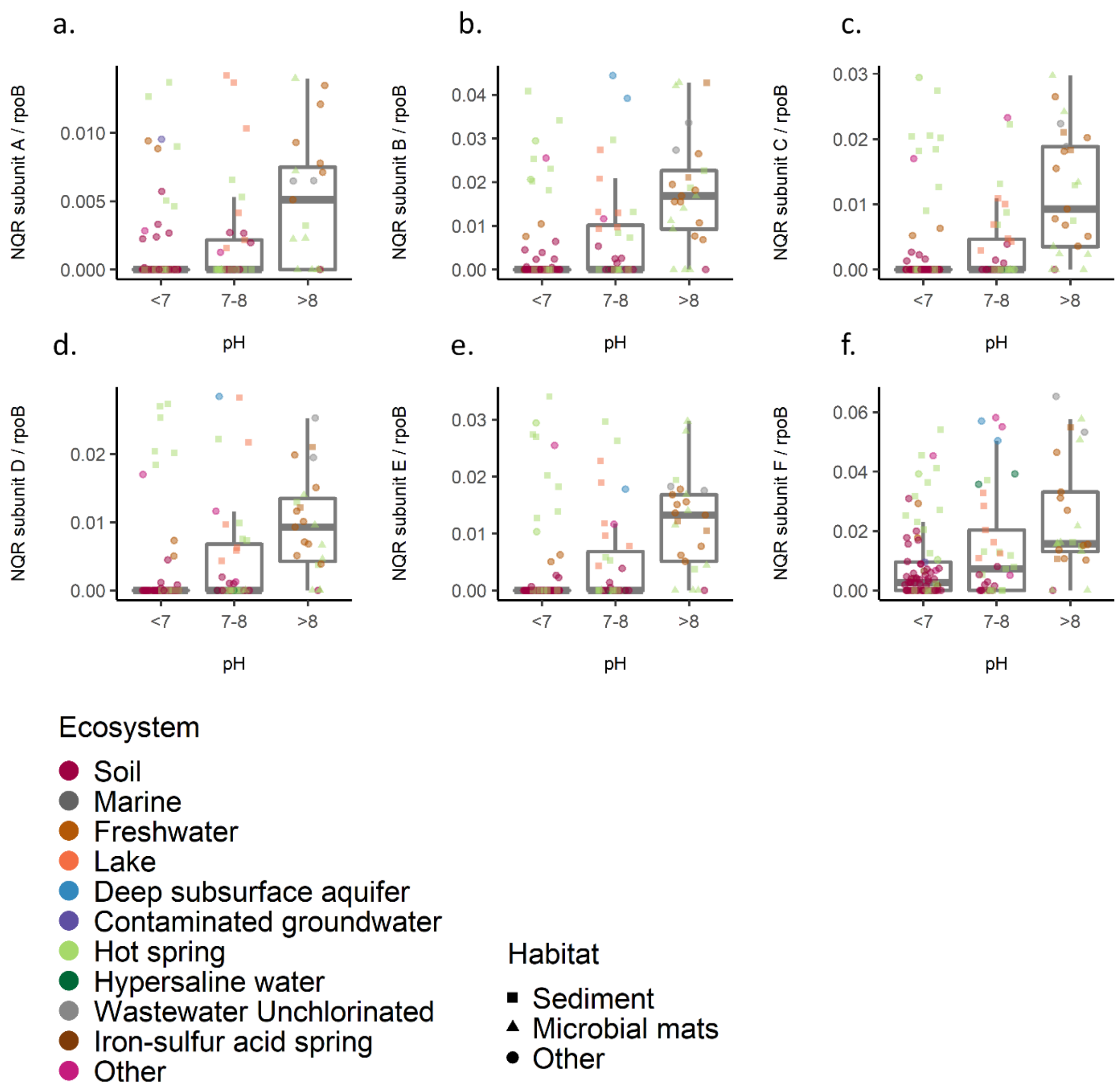


**Figure S6:** Meta- analysis of environmental metagenomes *nqr* gene count and pH concentration. Box plot of gene counts in acidic pH (<7), neutral (7-8), or alkaline (>8) pH for NQR operon (a-f subunits).

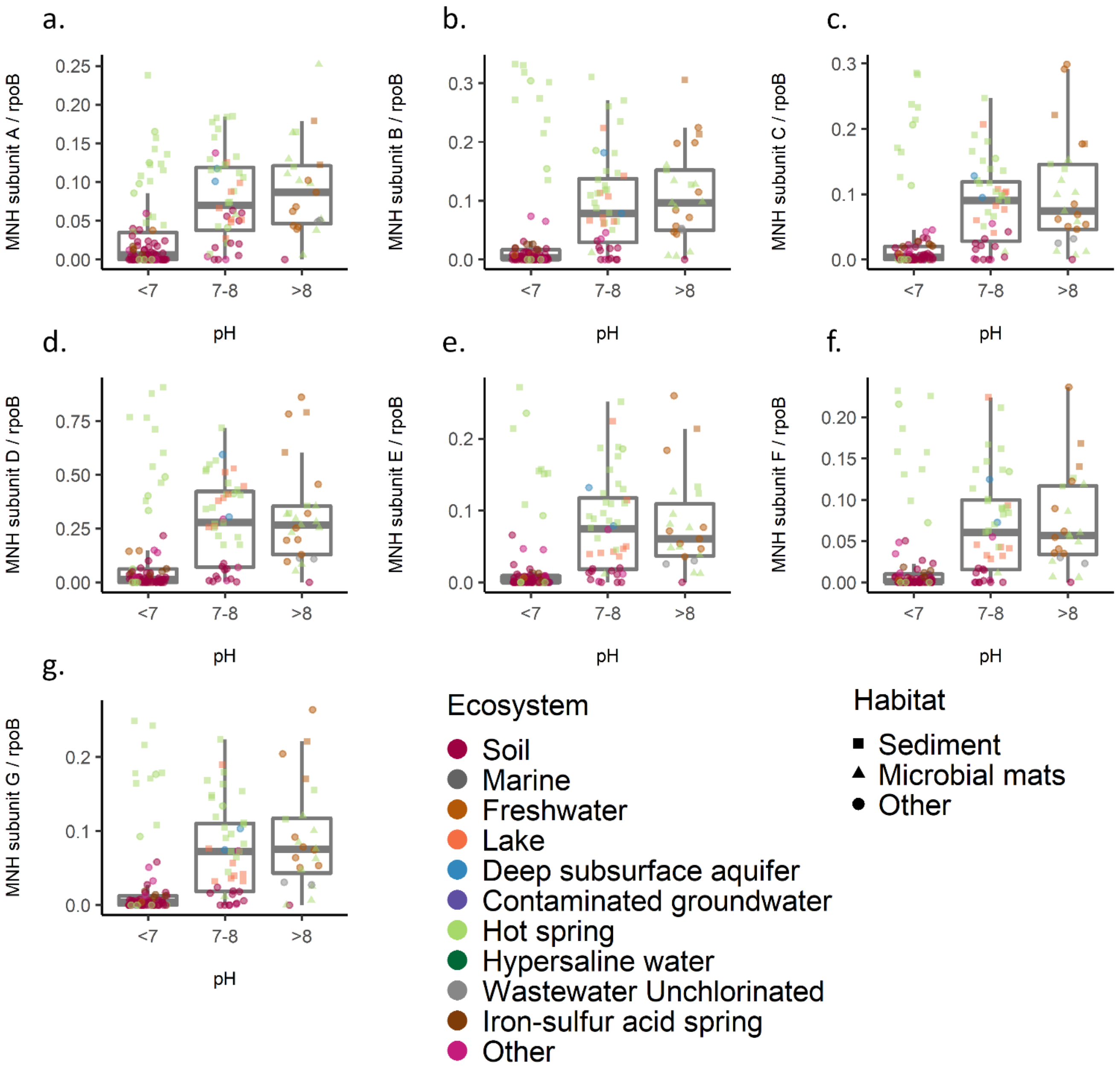


**Figure S7:** Meta- analysis of environmental metagenomes *mnh* gene count and pH concentration. Box plot of gene counts in acidic pH (<7), neutral (7-8), or alkaline (>8) pH for NQR operon (a-g subunits).

### Supplementary tables:

- **Table S1:** Detailed sequencing data: number of reads, contigs, N(50), percentage of mapped reads
- **Table S2:** Plant variables: yield, leaves cation concentration.
- **Table S3:** Plant RNAseq detailed EdgeR table.
- **Table S4:** Taxonomic affiliation: taxonomy detailed table. Including: count table, TMM normalized counts table, Deseq2.
- **Table S5: :** Tree file, based on MEGAN6 LCA algorithm
- **Table S6** Soil environmental variables: Na, K+, pH, EC, DOC, TN,
- **Table S7:** theSEED metagenome (DNA based) detailed table. Including: count table, TMM normalized table, DEseq2 result, goseq enrichment analysis.
- **Table S8:**Correlation of enriched SEED subsystems to environmental variables (pH, DOC, Na+, K+, DN)
- **Table S9:** KEGG metagenome (DNA based) detailed table. Including: count table, TMM normalized table, DEseq2 result, pathway and module enrichment analysis
- **Table S10:** Meta-analysis: JGI detailed table- projects id, habitat, pH concentration, oxygen concentration, gene counts
- **Table S11:** theSEED metatranscriptome (RNA based) detailed table. Including: count table, TMM normalized table, DEseq2 result, goseq enrichment analysis.
- **Table S12:** KEGG metatranscriptome (RNA based) detailed table. Including: count table, TMM normalized table, DEseq2 result, pathway and module enrichment analysis
